# Genome-wide meta-analysis of cognitive empathy: heritability, and correlates with sex, neuropsychiatric conditions and brain anatomy

**DOI:** 10.1101/081844

**Authors:** Varun Warrier, Katrina Grasby, Florina Uzefovsky, Roberto Toro, Paula Smith, Bhismadev Chakrabarti, Jyoti Khadake, Eleanor Mawbey-Adamson, Nadia Litterman, Jouke-Jan Hottenga, Gitta Lubke, Dorret I Boomsma, Nicholas G Martin, Peter K Hatemi, Sarah E Medland, David A Hinds, Thomas Bourgeron, Simon Baron-Cohen

## Abstract

We conducted a genome-wide meta-analysis of cognitive empathy using the ‘Reading the Mind in the Eyes’ Test (Eyes Test) in 88,056 research volunteers of European Ancestry (44,574 females and 43,482 males) from 23andMe Inc., and an additional 1,497 research volunteers of European Ancestry (891 females and 606 males) from the Brisbane Longitudinal Twin Study (BLTS). We confirmed a female advantage on the Eyes Test (Cohen’s d = 0.21, P < 2.2x10^−16^), and identified a locus in 3p26.1 that is associated with scores on the Eyes Test in females (rs7641347, P_meta_ = 1.58 x 10^−8^). Common single nucleotide polymorphisms (SNPs) explained 5.8% (95% CI: 0.45 - 0.72; P = 1.00 x 10^−17^) of the total trait variance in both sexes, and we identified a twin heritability of 0.28 (95% CI: 0.13-0.42). Finally, we identified significant genetic correlation between the Eyes Test and anorexia nervosa, measures of empathy (the Empathy Quotient), openness (NEO-Five Factor Inventory), and different measures of educational attainment and cognitive aptitude, and show that the genetic determinants of volumes of the dorsal striatum (caudate nucleus and putamen) are positively correlated with the genetic determinants of performance on the Eyes Test.

## Introduction

Cognitive empathy, defined as the ability to recognize what another person is thinking or feeling, and to predict their behaviour based on their mental states, is vital for interpersonal relationships, which in turn is a key contributor of one’s wellbeing. Cognitive empathy is distinct from affective empathy, defined as the drive to respond to another’s mental states with an appropriate emotion^1,2^. Difficulties in cognitive empathy have been found in different psychiatric conditions, particularly autism^3^. The dissociation between cognitive and affective empathy (the latter often being intact in autism, for example) suggests these have independent biological mechanisms.

Differences in cognitive empathy have been identified in individuals with different psychiatric conditions including autism^4^, schizophrenia^5,6^, and anorexia nervosa^7^. This includes both higher and lower cognitive empathy in comparison to neurotypical controls i.e. not empathizing enough, or empathizing too much, both of which can contribute to difficulties in social interactions and wellbeing^8^. However, little is known about the genetic correlates of cognitive empathy. It is unclear to what extent, genetically, differences in cognitive empathy are a risk factor for developing various psychiatric conditions. Further, as previous studies have typically been conducted using self-report or performance-based tests, results from these studies may be influenced by the characteristics of the test and/or factors associated with the psychiatric conditions themselves. In other words, from previous studies, it is difficult to tease apart the genetic and non-genetic contributions to strengths and difficulties in cognitive empathy, and how these relate to various psychiatric conditions.

Here, we investigated the genetic architecture of this aspect of social cognition using a well-validated test, the ‘Reading the Mind in the Eyes’ Test (Eyes Test). The Eyes Test is a brief online test where participants are shown photographs of the eye regions and have to identify the appropriate emotion or mental state they express^2^. It has been widely used to investigate differences in cognitive empathy in several neuropsychiatric conditions including autism^4^, schizophrenia^9^, bipolar disorder^10^, anorexia nervosa^11^, and major depressive disorder^12^. We conducted a genome-wide association meta-analysis of cognitive empathy in more than 89,000 individuals of European ancestry, and investigated both SNP-based and twin-based heritabilities. We further conducted bivariate genetic regression analyses for psychiatric conditions, psychological traits, and brain volumes. We finally conducted gene based enrichment analysis and investigate potential genetic sources of sex differences.

## Methods

### Participants

#### 23andMe

Research participants were customers of 23andMe, Inc., and have been described in detail elsewhere^13,14^. All participants completed an online version of the ‘Reading the Mind in the Eyes’ test (Eyes Test)^2^ online on the 23andMe research participant website (36 items). In total, 88,056 participants (44,574 females and 43,482 males) of European ancestry completed the Eyes Test and were genotyped. All participants provided informed consent and answered questions online according to 23andMe’s human subjects protocol, which was reviewed and approved by Ethical & Independent Review Services, an AAHRPP-accredited private institutional review board (http://www.eandireview.com). Only participants who were primarily of European ancestry (97% European Ancestry) were selected for the analysis using existing methods^15^. Unrelated individuals were selected using a segmental identity-by-descent algorithm^16^.

#### Brisbane Longitudinal Twin Study (BLTS)

In addition, 1,497 participants (891 females and 606 males) of Caucasian ancestry with genotype data from the BLTS completed the short version (14 questions) of the Eyes Test online as part of a study on genetic and environmental foundations of political and economic behaviors^17^. Participant ages ranged from 18 to 73 (M = 37, *SD* = 14). All participants provided online consent and the study was approved by the QIMR Berghofer Human Research Ethics Committee.

### Measures

The ‘Reading the Mind in the Eyes’ Test is a brief questionnaire of cognitive empathy. Participants are shown scaled, black and white photographs of eye regions of actors and they have to choose the cognitive state portrayed from the four options provided. The Eyes Test has good test-retest reliability (eg: reliability of 0.833 in the Italian version^18^, and 0.63 in the Spanish version^19^)^4^, and scores are unimodally and near-normally distributed in the general population. In the BLTS dataset, there was a modest test-retest correlation of 0.47 in 259 participants who retook the test after a gap of nearly two years (Supplementary Note section 1). For each correct answer on the Eyes Test, participants score 1 point, so the scores ranged from 0 - 36 on the full version of the Eyes Test and 0 - 14 on the short version of the Eyes Test. Further details are provided in the Supplementary Note section 1.

### Genotyping, imputation and quality control

#### 23andMe cohort

DNA extraction, genotyping, imputation and initial quality control were completed by 23andMe, Inc. Participants provided saliva samples, and DNA extraction and genotyping were performed by the National Genetic Institute. All participants were genotyped using one of four different platforms (V1, V2, V3 and V4). Briefly, the V1 and V2 chips were based on the Illumina Human Hap550+ BeadChip (560,000 SNPs), the V3 on the Illumina OmniExpress+ Beadchip (950,000 SNPs). The V4 had a fully customized array of approximately 570,000 SNPs. Across all platforms, a total of 1,030,430 SNPs were genotyped. For this analysis, we included only participants with a call rate greater than 98.5, and SNPs that passed the Hardy-Weinberg Equilibrium Test at P < 10^−20^ and had a genotype rate > 90%. In addition, SNPs present only on platform V1, Chromosome Y and mitochondrial SNPs were excluded due to small sample sizes and unreliable genotype calling respectively. Using trio-data, where available, SNPs that failed the parent-offspring transmission test were also excluded. Imputation was performed using Minimac2^20^ using the September 2013 release of the 1000 Genomes Phase 1 reference haplotypes phased using Beagle4^21^ (V3.3.1). Our analyses were restricted to SNPs that had a minor allele frequency of at least 1%, which left 9,955,952 SNPs after quality control. Genotyping, imputation, and preliminary quality control were performed by 23andMe.

#### BLTS cohort

The BLTS participants were genotyped on Illumina Human610-Quadv1_B or HumanCoreExome-12v1-0_C chips. These samples were genotyped in the context of a larger genome-wide association project. Genotype data was screened for genotyping quality (GenCall < 0.7 from the Human610-Quadv1_B chip), individual and SNP call rates (< 0.95 and < 0.99 for exome markers on the HumanCoreExome-12v1-0_C chip), Hardy-Weinberg Equilibrium (P < 10^−6^), and MAF (< 0.01). The data were checked for non-European ancestry, pedigree, sex, and Mendelian errors. Data from the two different chips were separately phased using SHAPEIT2 and imputed to the 1000 Genomes reference panel (Phase 1 v3) using Minimac3. After imputation SNPs with a MAF < 0.05% were excluded, leaving 11,133,794 SNPs for analyses. We further excluded SNPs with imputation r^2^ < 0.6 for meta-analysis.

### Statistical analyses

#### Association analyses

Linear regression for the 23andMe cohort was performed for the Eyes Test scores using age, sex, and the first four ancestry principal components as covariates. For the sex-stratified analyses, sex was excluded as a covariate. The same regression model was used for the BLTS after accounting for relatedness using RAREMETALWORKER. Inverse variance weighted meta-analysis was performed using Metal^22^. Post meta-analysis, we excluded SNPs that were only genotyped in the BLTS cohort due to the small sample size, but included SNPs that were only genotyped in the 23andMe cohort. LD pruning was performed using Plink^23^ with an r^2^ of 0.1. We calculated the variance explained by each individual SNP^24^ using the following formula:

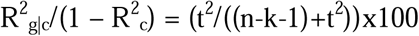

Where R^2^_g|c_/(1 - R^2^_c_) is the proportion of variance explained by the SNP after accounting for the effects of the covariates, t is the t-statistic of the regression co-efficient, k is the number of covariates, and n is the sample size. We corrected for winner’s curse using an FDR based approach^25^.

#### Heritability and genetic correlation

We used Linkage Disequilibrium Score regression coefficient (LDSR to calculate genomic inflation in the meta-analysis due to population stratification^26^ (https://github.com/bulik/ldsc). Intercepts for the non-stratified GWAS was 1.01 (0.006), for the males-only GWAS was 1.006 (0.006), and for the females-only GWAS was 1.005 (0.006). SNP heritability and genetic correlation were calculated using LDSR. Difference in heritability between males and females was quantified using:

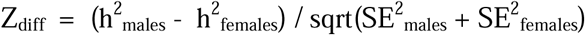

Where Z_diff_ is the Z score for the difference in heritability for a trait, (h^2^_males_ - h^2^_females_) is the difference SNP heritability estimate in males and females, and SE is the standard errors for the heritabilities. We calculated two-tailed P-values in R. We performed genetic correlation using summary GWAS data using LDSR. For all genetic correlation analyses, we used LD data from the North West European population as implemented in LDSR. Intercepts were not constrained in the analyses. We used a Benjamini-Hochberg FDR approach and a q-value threshold of 0.05 for determining statistical significance of the genetic correlations. Further details about the samples used are provided in the Supplementary Note section 2.

#### Twin Heritability

Twin heritability was estimated from 749 twin individuals (including 122 complete monozygotic pairs and 176 complete dizygotic pairs) in the BLTS using full information maximum likelihood in OpenMx^27^ in R, which makes use of all available data. All twins completed the short version of the Eyes Test, and for those who completed the test twice only their first attempt was included in analyses. ADE, ACE, AE, CE, and E models were fit to the data and fit indices compared to determine the best-fitting model. Standardised variance components are reported from the best-fitting model, the AE model. Further details are given in the Supplementary Note section 4.

#### Gene-based analyses and sex difference analyses

We used MetaXcan^28^ using tissue weights from the GTEx to perform gene-based analysis (https://github.com/hakyimlab/MetaXcan). MetaXcan uses summary statistics to perform gene based association analyses. It incorporates eQTL data from the GTEx consortium to infer gene level expression based on the summary GWAS statistics provided. This can be used to identify tissue-specific gene expression for the trait of interest. Here, we performed gene based analysis for the non-stratified GWAS metaanalysis for nine neural tissues: anterior cingulate cortex (BA24), caudate basal ganglia, cerebellar hemisphere, cerebellum, cortex, frontal cortex (BA9), hippocampus, hypothalamus, nucleus accumbens basal ganglia, and putamen basal ganglia, using gene-expression regression co-efficients for these tissues from the GTEx project. This is based on tissues from 73 - 103 individuals. We chose neural tissues as we hypothesized that cognitive empathy is largely a neural phenotype. As MetaXcan predicts expression level from SNP information, we filtered out genes whose correlation with predicted models of expression was < 0.01, as incorporated in MetaXcan. This steps helps guard against false positives, by removing genes whose expressions are poorly predicted by the model. We used an FDR based correction to correct for all the tests run across all the tissues. Details of sex-difference analyses are provided in Supplementary Note section 3.

## Data Availability

Summary level data may be requested from 23andMe, Inc. and received subject to 23andMe's standard data transfer agreement. We have also provided summary statistics for the first 10,000 LD pruned SNPS for the three GWAS analyses (males-only, females-only, and non-stratified) as supplementary data.

## Results

### Heritability

In collaboration with 23andMe, Inc. and the Brisbane Longitudinal Twin Study (BLTS) cohort, we conducted three separate genome-wide association study meta-analyses(GWASMAs) of the Eyes Test: a males-only GWAS (n = 43,482), a females-only GWAS (n = 44,574), and a non-stratified GWAS (n = 88,056). The study protocol is provided in Figure 1. All participants from the 23andMe cohort completed the full version of the Eyes Test online, comprising 36 questions (mean score = 27.47±3.67), while participants from the BLTS cohort completed the short version of the Eyes Test (14 questions, mean = 8.85±2.34) (Supplementary Note Section 1). Scores on the Eyes Test were significantly associated with age and sex in the 23andMe cohort (age: −0.026±0.0007; P < 2.2x10^−16^, sex (females): 0.77±0.02; P < 2.2x10^−16^). We used LDSR to calculate the heritability explained by all the SNPs in the HapMap3 with minor allele frequency > 5%. We identified a significant narrow sense heritability of 5.8% (95% CI: 4.5% - 7.2%; P = 1.00 x 10^−17^) in the non-stratified GWAS. We calculated the twin heritability from 749 twin individuals (including 122 complete monozygotic pairs and 176 complete dizygotic pairs) in the BLTS. Heritability, from the best-fitting additive genes/unique environment (AE) model, was 0.28 (95% CI: 0.13-0.42) (Supplementary Note Section 4).

**Figure 1:**
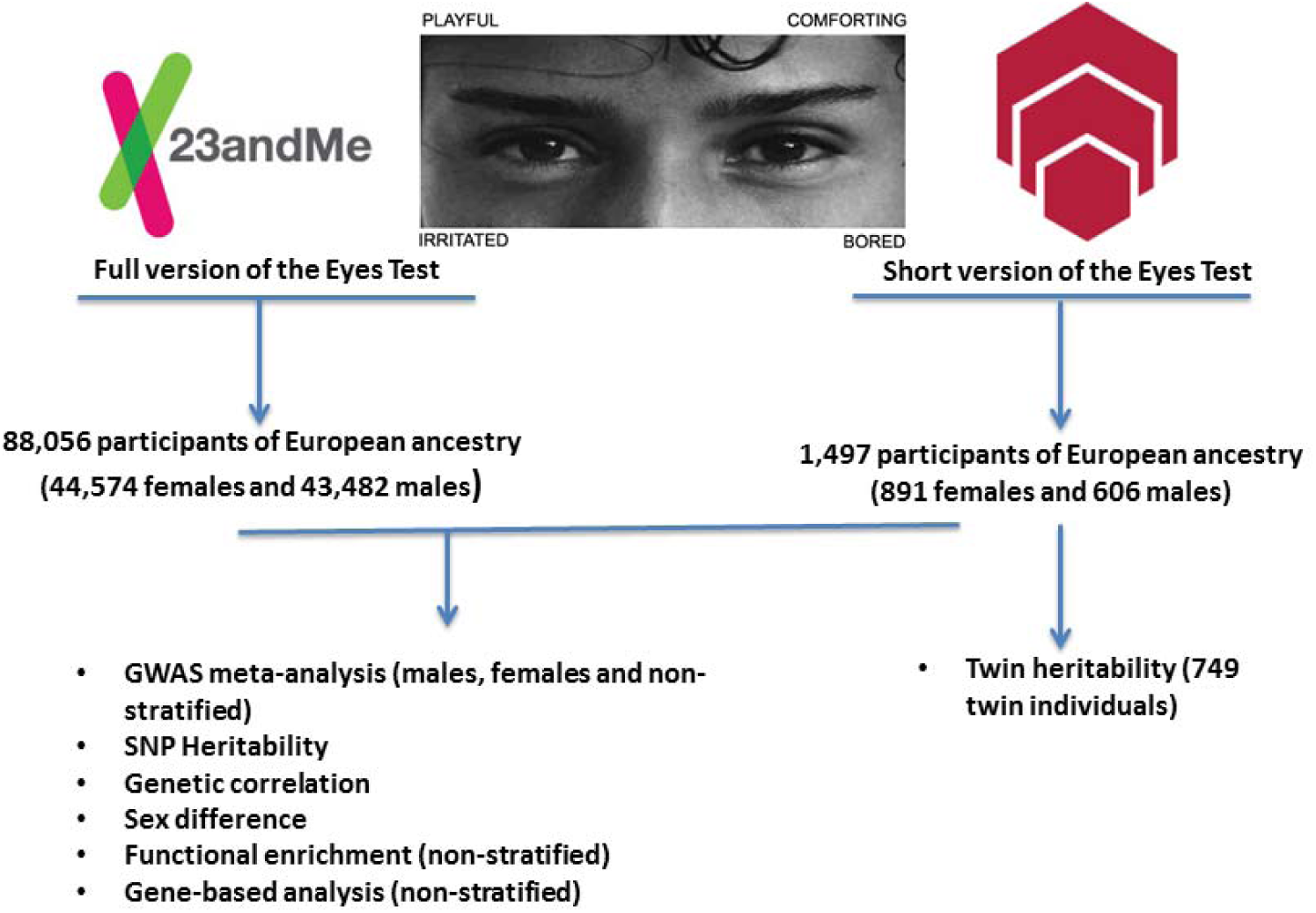
Schematic diagram of the study protocol. 88,056 Caucasian participants from 23andMe, Inc. completed the full version of the Eyes Test and were genotyped. An additional 1,497 Caucasian participants from the Brisbane Longitudinal Twin Study completed the short version (14 questions) of the Eyes Test and genotyped. Genome-wide association meta-analysis was performed on the combined cohort of 89553 participants. Three separate meta-analyses were performed: males-only, females-only, and non-stratified. Subsequently, functional enrichment and gene-based analysis was performed for the non-stratified meta-analysis GWAS using the 23andMe dataset. SNP heritability and genetic correlation using LDSR was performed for the 23andMe GWAS dataset. Sex differences were also investigated using the same dataset. In parallel, twin heritability was calculated from 749 twin individuals from the Brisbane Longitudinal Twin Study who had completed the short version of the Eyes Test.

### Genetic correlation

We next investigated how the non-stratified Eyes Test is genetically correlated to psychiatric conditions and specific psychological and cognitive traits for which summary GWAS data were available (Supplementary Table 5). After correcting for multiple testing, we identified significant positive genetic correlations between Eyes Test scores and two cognitive traits: self-reported empathy measured using the Empathy Quotient (r_g_ = 0.18±0.07; P__FDRadjusted_= 0.031)^29^, and the NEO-Five Factor Inventory measure of openness (r_g_ = 0.54±0.14; P__FDRadjusted_= 9.73x10^−4^)^30^. We also identified significant positive correlations with different measures of cognition and education: college years (r_g_ = 0.40±0.6; P__FDRadjusted_ = 3.1x10^−10^)^31^, educational attainment (0.34±0.04; P__FDRadjusted_=3.7x10^−16^)^32^, and childhood cognitive aptitude (calculated as Spearman’s g and is, hence, independent of word knowledge)^33^ (r_g_ = 0.34±10; P__FDRdjusted_ =0.0075). In addition, we identified a significant positive genetic correlation between the Eyes Test scores and anorexia nervosa (Anorexia-GCAN r_g_ = 0.14±0.06; P__FDRadjusted_ = 0.047), which we validated using a non-independent data-freeze with more cases for anorexia nervosa from the Psychiatric Genomics Consortium (Anorexia-PGC r_g_ = 0.25±0.08; P__FDRadjusted_ = 0.009) (Figure 2). We did not identify a significant genetic correlation between autism and scores on the Eyes Test.

We also investigated if subcortical brain volumes are correlated with performance on the Eyes Test. We used data from the ENIGMA consortium for six subcortical regions and intracranial volume^24^. We excluded the amygdala, even though it is relevant for social cognition, as the low heritability of the amygdala could not be accurately quantified using LDSR^23^. After correcting for multiple testing, we identified a significant positive correlation between the Eyes Test scores and the volumes of the caudate nucleus^24^ (r_g_ = 0.24±0.09; P__FDRadjusted_= 0.033) and volume of the putamen (r_g_ = 0.21±0.08; P__FDRadjusted_ = 0.041), which together form the dorsal striatum. All genetic correlations are provided in Supplementary Table 5.

We also investigated sex-stratified genetic correlations between the Eyes Test and educational attainment, the only relevant phenotype where we had access to sex-stratified data. We identified a modest, significant genetic correlation between educational attainment and the Eyes Test in the males-only dataset: r_g_ = 0.23±0.05; P = 2.6x10^−5^. We identified a higher, significant genetic correlation between educational attainment and Eyes Test in the females-only dataset: r_g_ = 0.39±0.06; P = 5.88x10^−11^. These results suggest that females share greater pleiotropy between general cognition and cognitive empathy than males, indicating different genetic mechanisms for the development of cognitive empathy.

**Figure 2:**
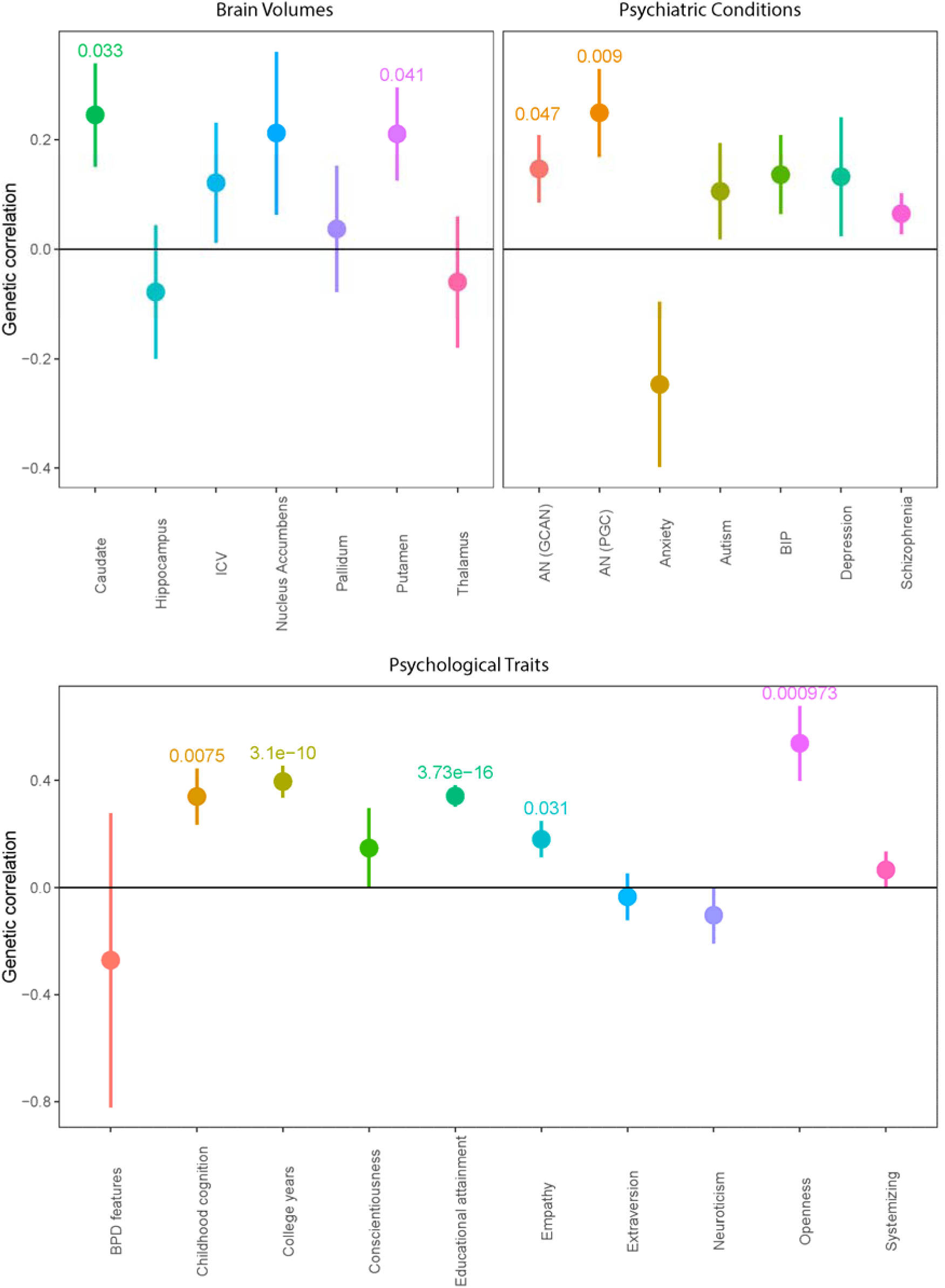
Genetic correlations between the Eyes Test and psychiatric conditions, psychological traits and subcortical brain volumes. Genetic correlations and standard errors for the Eyes Test in the 23andMe cohort. Figures above the bars represent FDR adjusted P-values. All P-values with P _FDRcorrected_ < 0.05 provided. * represents significant P-values after FDR correction. Point estimate represents the genetic correlation, and the error bars represent the standard errors. AN (GCAN) is the Anorexia Nervosa-GCAN dataset, AN (PGC) is the Anorexia Nervosa PGC dataset, BIP is bipolar disorder, BPD features is borderline personality disorder features, ICV is intracranial volume. We have removed the genetic correlation for agreeableness from this figure due to the high standard errors. The genetic correlations, standard errors, and P-values for all traits including agreeableness are provided in Supplementary Table 5.

### Genome-wide association meta-analyses

GWAMA of the non-stratified and the males-only datasets did not identify any significant loci. In the females-only analysis, we identified one locus at 3p26.2 that was significant at a threshold of P < 5x10^−8^. This locus contains 21 significant SNPs in high LD with the leading SNP rs7641347 (Figure 2), with concordant effect direction for 19 SNPs in the 23andMe and BLTS datasets. The leading SNP rs7641347 (P_meta_ = 1.58 x 10^−8^) explained 0.067% of the total variance, or 0.013% of the total variance after correcting for winner’s curse^25^. Of the two SNPs with discordant effects in the two datasets, rs114076548 was the most-significant SNP in the 23andMe dataset and had P = 6.49x10^−9^. We did not identify any inflation in the P-values of the GWAMA due to population stratification using LDSR (intercept = 1.01 ± 0.007).

**Figure 3:**
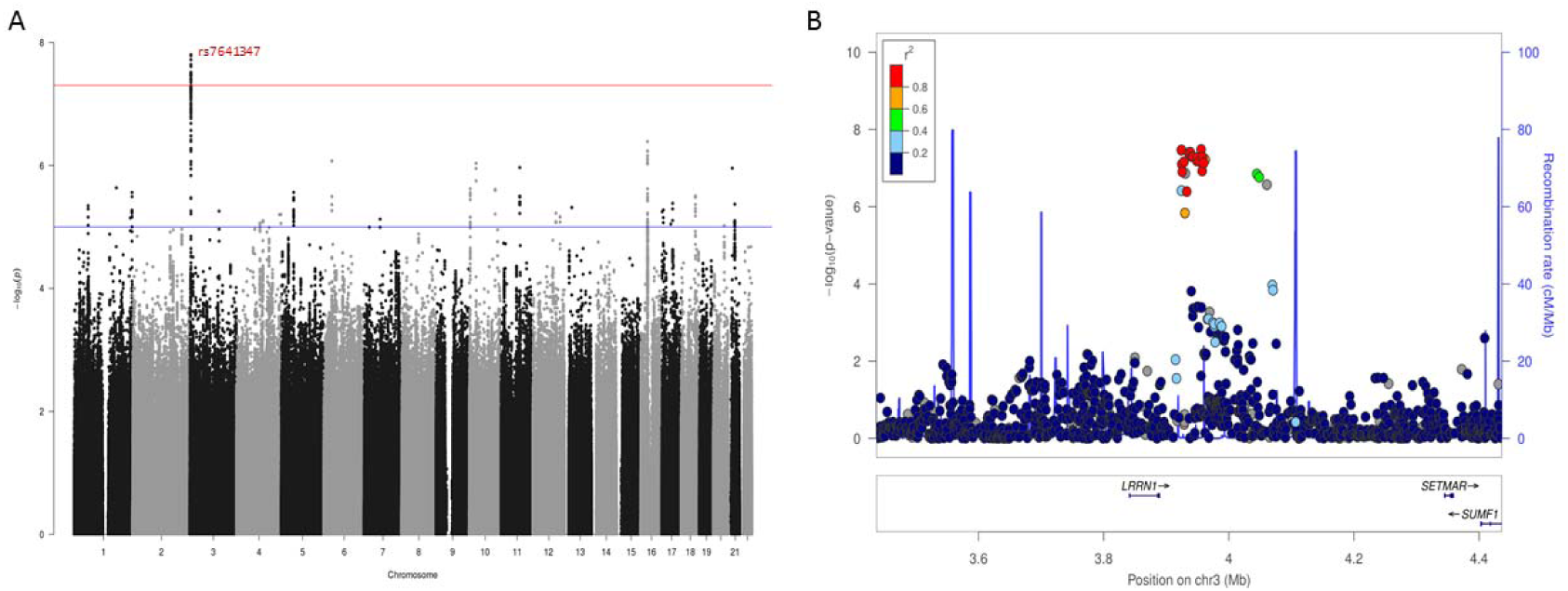
Manhattan plot and regional association plot for the Eyes Test (females) meta-analysis GWAS. A. Manhattan plot of the Eyes Test meta-analysis (female). X axis is the chromosomal position of the SNP, and Y axis is the negative logarithm of the P-value. The red line indicates genome-wide significant threshold of 5×10^−8^. Lead SNP for all loci with P < 1×10^−6^ is provided. n = 44,574, and λ_gc_ = 1.05. LDSR intercept = 1.05. Regional association plot of the significant locus for the Eyes Test (females) meta-analysis.

The leading SNP (rs7641347) is located in an intron of *SUMF1* and was nominally significant in the non-stratified analysis (P_meta_ = 1.1x10^−5^), but non-significant in the males-only analysis (P_meta_ = 0.4954). In addition, SNPs in high LD (r^2^ > 0.8) were also not nominally significant in the males-only analysis. Together, all 21 SNPs span a region of approximately 77kb 3p26.2 (Supplementary Table 1). At this locus, in addition to *SUMF1,* 2 other genes are present: Leucine Rich Neuronal 1 *(LRRN1)* and SET Domain And Mariner Transposase Fusion Gene *(SETMAR). LRRN1* is highly expressed in brain tissues^34^, with median expression the highest in the putamen, nucleus accumbens and the caudate nucleus, all three of which are part of the striatum. Deletion of 3p26.1 and 3p26.2 can cause developmental delay, hypotonia and epileptic seizures and has been implicated in autism^35^.

The most significant SNP in the males-only GWAS meta-analysis (rs4300633 in 16p12.3, P = 9.11x10^−8^) explained 0.062% of the variance, and the most significant SNP in the non-stratified GWAS meta-analysis (rs149662397 in 17q21.32 P = 1.58x10^−7^) explained only 0.029% of the variance. All LD pruned SNPs with P < 1x10^−6^ are provided in Supplementary Table 2. The QQ-plot and locus-zoom plot for the females-only meta-analysis, and the Manhattan and QQ-plots for the males-only and non-stratified analyses are provided in the Supplementary Note section 5. Gene-based analyses MetaXcan^28^ for ten neural tissues (Methods) and functional enrichment analyses for the non-stratified GWAS did not identify any significant results (Supplementary Tables 3 and 4).

### Sex differences

We also investigated sex-differences in the Eyes Test. There was a significant female advantage on the scores of the full Eyes Test (males = 27.08±3.75; females = 27.85±3.55; cohen’s d = 0.21, P < 2x10^−16)^’ replicating previous results^36^ (Figure 4). There was no significant difference in males-only or females-only SNP heritability estimates (males = 0.071±0.011, females = 0.067±0.011; P = 0.79). There was a reasonably high but incomplete genetic correlation between males and females (r_g_ = 0.68±0.12; P = 2.70x10^−8^). Binomial sign test of LD-pruned nominally significant SNPs in the sex-stratified analyses identified that 61% (95% CI: 59% - 62%) of the SNPs had a concordant effect direction (P < 2.2x10^−16^). We further investigated the effect direction and statistical significance of all independent SNPs with P < 1x10^−6^. SNPs that were of suggestive significance in one sex were not nominally significant in the other (Supplementary Table 2 and Supplementary Note section 5). Using MetaXcan^28^ we identified the top cortically expressed genes (P < 0.05) for both sexes and calculated the overlap in the genes. We did not find any enrichment in gene overlap (Fold difference = 1.2, P = 0.264). We also investigated if there was an enrichment of female-overexpressed or male-overexpressed cortical genes for the Eyes Test (Methods) and did not find any significant enrichment (Supplementary Note section 3, and Supplementary Tables 6-8).

**Figure 4.**
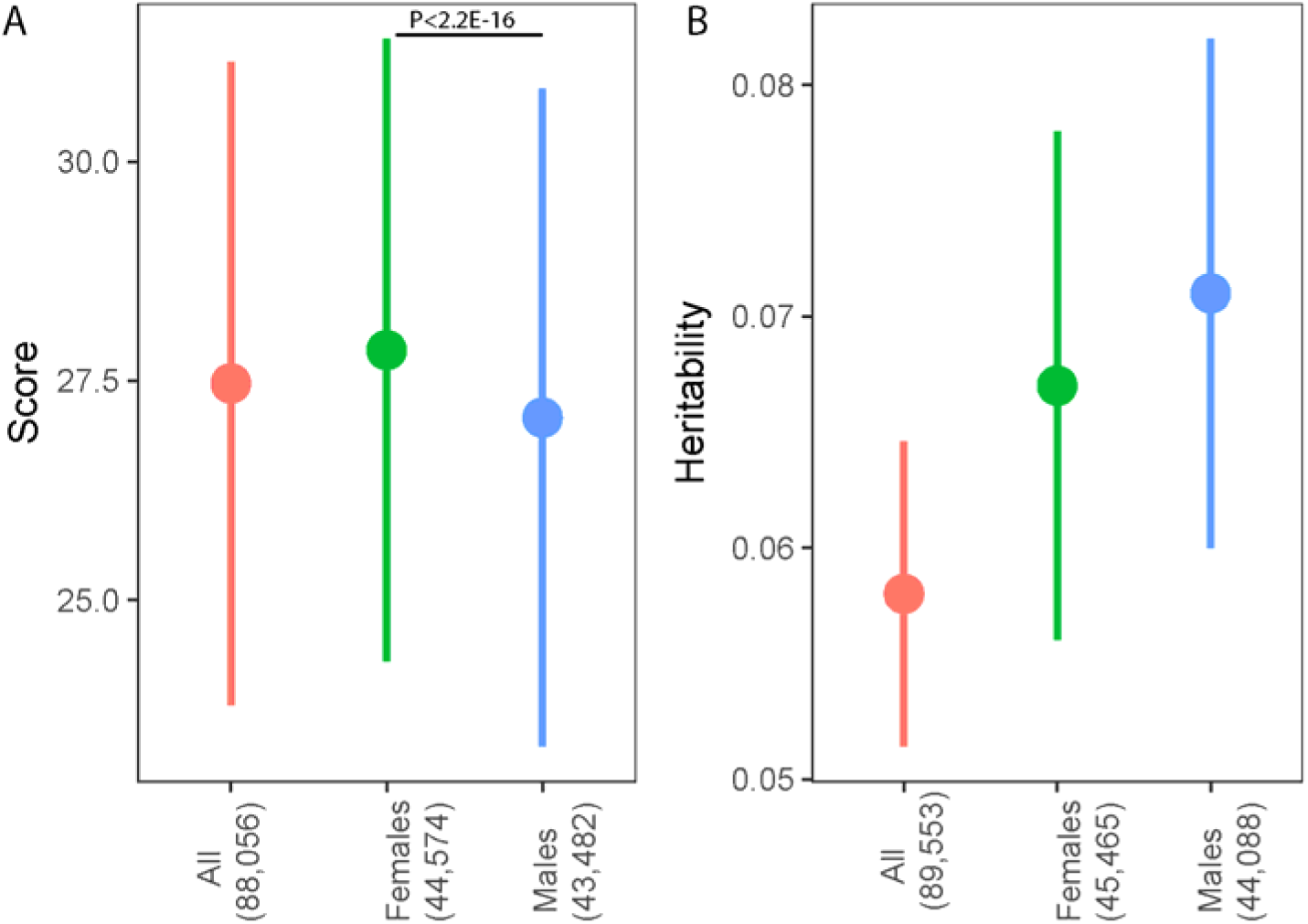
Mean scores and SNP heritability. 4A. Mean phenotypic scores and standard deviations for the Eyes Test in the 23andMe cohort. Point estimate provides the mean score, and the error bars represent standard deviations. Difference in mean scores between males and females was highly significant (P < 2.2E-16; Cohen’s d = 0.21). Numbers in brackets indicate the number of participants in each GWAS. All: non-stratified GWAS; Females: Females-only GWAS; Males: Males-only GWAS. 4B. Mean SNP heritability estimates and standard errors for the Eyes Test in the GWAMA. Point estimate provides mean SNP heritability, and error bar represents standard errors. There was no significant difference in SNP heritability estimates between males and females (P = 0.79). Numbers in brackets indicate the number of participants in each GWAMA. All: non-stratified GWAMA; Females: Females-only GWAMA; Males: Males-only GWAMA

## Discussion

This is the first large-scale genetic study investigating the genetic architecture of cognitive empathy. We investigated heritability estimates of the Eye Test. In our sample of 749 twin individuals, heritability was approximately 0.28 (95% CI: 0.13-0.42). This is in keeping with previous studies that have investigated heritability of other facets of empathy in twins. A meta-analysis of empathy in twins identified that approximately a third of the variance is heritable^37^. In our sample of 88,056 unrelated research volunteers from 23andMe Inc, SNP-based heritability was estimated using LDSR, and approximately 5% of the trait was additively heritable. It is likely that heritability of cognitive empathy changes with age (which was significantly correlated with scores on the Eyes Test in this dataset), as is observed in prosocial behaviour^37^. In our analyses, age was included as a covariate, and thus our SNP heritability is likely to represent the lower bound of the SNP heritability.

We identified significant genetic correlation between volume of the caudate nucleus, putamen and scores on the Eyes Test. Imaging studies have reported activation in both the putamen^38^ and caudate nucleus^39^ on tasks of social cognition. In humans, the ventral striatum is composed of the nucleus accumbens and olfactory tubercle, whereas the dorsal striatum is composed of the caudate nucleus and putamen. There is some evidence to support the role of the striatum in theory of mind^40^, though there is no clear consensus that cognitive and affective empathy utilize different neural circuits. Our results suggest that variants that contribute to volumes of regions in the striatum, the dorsal striatum in particular, also contribute to cognitive empathy. It is difficult to delineate the mechanism behind this observed pleiotropy, but we can hypothesize that volumes of striatal structures influence scores on the Eyes Test.

In addition, we identified significant positive genetic correlations with different measures of cognitive ability including educational attainment. This reflects phenotypic correlation between measures of cognitive empathy and cognitive ability. A meta-analysis has identified a significant positive correlation between scores on the Eyes Test and IQ (n = 3583; r = 0.24; 95% CI: 0.16 - 0.32)^41^. Other tests of theory of mind are also positively correlated with cognitive aptitude and measures of intelligence^42–44^. We also found a significant positive genetic correlation with the NEO-Openess to experience, and self-reported empathy measured using the Empathy Quotient. Again, both the genetic correlations reflect previous correlation at a phenotypic level between measures of empathy and personality^45^. With psychiatric conditions, there was a significant positive correlation with anorexia which we identified using two non-independent dataset. One study has identified that individuals with anorexia report higher personal distress^46^, though other studies have reported that deficits in social cognition may be attributable to comorbid alexithymia^47^. Our research suggests that higher genetic contribution to cognitive empathy increases the genetic risk for anorexia. However, it is likely that the development of anorexia can contribute to poorer performance on tests of social cognition and cognitive empathy through other factors including alexithymia.

We did not identify a significant genetic correlation between the Eyes Test and autism. Recent research suggests that this may be due to heterogeneity in performance in the Eyes Test, with only a subset of individuals with autism showing impaired performance on the Eyes Test^4^. In addition, the cognitive phenotype of autism involves non-social aspects (such as attention to detail), not just social ones. A meta-analysis has reported global or selective deficits in performance on the Eyes Test in individuals with schizophrenia, anorexia, bipolar disorder, and clinical depression but preserved or even enhanced performance for individuals with non-clinical depression and borderline personality disorder^48^. However, these studies are typically conducted in small sample sizes and presence of psychiatric conditions may impair performance on the Eyes Test. Further, performance on the Eyes Test may be influenced by multiple factors related to psychiatric conditions, and need not necessarily measure the causal relationship between psychiatric conditions and cognitive empathy.

We also identified one locus that is significantly associated with empathy in females. The top SNP (rs7641347) had a P-value = 1.58x10^−8^. One of the closest gene, *LRRN1,* is highly expressed in striatum according to the GTEx database. However, we were unable to identify any eQTL that specifically linked this locus to the gene. *LRRN1* is a gene that is not well characterized. In chicks, Lrrn1 is necessary for the formation of the mid-brain hind-brain boundary^49^. The locus was significant in females, nominally significant in the non-stratified analyses, and non-significant in the males-only analyses, suggesting a sex specific involvement of this locus in cognitive empathy measured using the Eyes Test. We note that even with approximately 90,000 individuals this GWAMA was underpowered to detect loci of significant effect, owing to the relatively low variance explained per SNP.

It was also interesting to note that while twin and SNP-based heritability did not vary between the sexes in our study, we replicated the female-advantage on the Eyes Test in the largest sample thus far. Sex-stratified analyses also allowed us to investigate the genetic correlation between males and females, and subsequently, sex-specific imputed gene expression in cortical tissues. Male-female genetic correlation was only modest, which was supported by binomial sign test. In comparison, other traits for which we had sex-stratified data, genetic correlation was considerably higher (eg: self-reported empathy^29^: r_g_ = 0.82, sd =0.16, systemizing^29^: r_g_ = 1.0, sd = 0.16; educational attainment^32^: r_g_ = 0.91, sd = 0.02). Further, examining the top SNPs indicated a sex-specific architecture for the most significant set of SNPs (P < 1x10^−6^). We also did not identify a significant overlap between the genes identified for the sex-stratified GWAS. All of this suggests that there is some sex specificity in the genetic architecture of cognitive empathy. How this sex-specific architecture is expressed and interacts with steroid hormones will help shed further light on the biological contributions to the female superiority on the Eyes Test.

In conclusion, we identify a genetic locus that is associated with scores on the Eyes Test in females. We identify significant positive genetic correlations between scores on the Eyes Test and empathy, cognitive aptitude and educational attainment, and openness to experience. We also identify positive correlation between striatal volumes and scores on the Eyes Test. Phenotypic sex-differences for the Eyes Test may be partly due to different genetic architectures in males and females, interacting with postnatal social experience.

## Acknowledgements

This study was funded by grants from the Templeton World Charity Foundation, Inc., the Medical Research Council, the Wellcome Trust, the Autism Research Trust, the Institut Pasteur, the CNRS and the University Paris Diderot. VW is funded by St. John’s College, Cambridge, and Cambridge Commonwealth Trust. The research was carried out in association with the National Institute for Health Research (NIHR) Collaboration for Leadership in Applied Health Research and Care East of England at Cambridgeshire and Peterborough NHS Foundation Trust. The views expressed are those of the author(s) and not necessarily those of the NHS, the NIHR or the Department of Health. The research was supported by the National Institute for Health Research (NIHR) Collaboration for Leadership in Applied Health Research and Care East of England at Cambridgeshire and Peterborough NHS Foundation Trust and the National Human Genome Research Institute of the National Institutes of Health (grant number R44HG006981). The National Science Foundation (grant numbers 0729493 and 0721707) supported the research on the Brisbane Longitudinal Twin Study. FU was supported by the British Friends of Haifa University, the Israel Science Foundation (grant no. 449/14), the British Friends of Hebrew University, and the Joseph Levy Charitable Foundation. TB was supported by the Institut Pasteur, the Bettencourt-Schueller Foundation, University. We thank the research participants and employees of 23andMe for making this work possible. We also thank the volunteers of the Brisbane Longitudinal Twin Study and The NIHR Cambridge BioResource. Finally, we thank Carrie Allison, Michael Lombardo, Owen Parsons, and Richard Bethlehem for helpful discussions.

